# Interdependent Dynamics of mRNA Expression and HIV-1 Viral Load: Insights from Transcriptomics and Mendelian Randomization

**DOI:** 10.1101/2025.11.13.688267

**Authors:** Sergey Oreshkov, Christian W. Thorball, Jenny Maylan, Maude Muriset, Alexandra Calmy, Angela Ciuffi, Luigia Elzi, Johannes Nemeth, Ansar Muhammad, Lukas Baumann, Patrick Schmid, Marcel Stöckle, Daniel Alpern, the Swiss HIV Cohort Study, Jacques Fellay, Federico Santoni

**Affiliations:** Service of Endocrinology, Diabetology and Metabolism, Lausanne University Hospital, Lausanne, Switzerland; Faculty of Biology and Medicine, University of Lausanne, Lausanne, Switzerland; Biomedical Data Science Center, Lausanne University Hospital and University of Lausanne, Lausanne, Switzerland; School of Life Sciences, École Polytechnique Fédérale de Lausanne, Lausanne, Switzerland; Division of Infectious Diseases, Geneva University Hospitals, Geneva, Switzerland; Institute of Microbiology, Lausanne University Hospital and University of Lausanne, Lausanne, Switzerland; Division of Infectious Diseases, Regional Hospital Bellinzona, Bellinzona, Switzerland; Department of Infectious Diseases and Hospital Epidemiology, University Hospital Zurich, University of Zurich, Zurich, Switzerland; Department of Ophthalmology, University of Lausanne, Jules-Gonin Eye Hospital, Fondation Asile Des Aveugles, Lausanne, Switzerland; Department of Infectious Diseases, Inselspital Bern University Hospital, University of Bern, Bern, Switzerland; Division of Infectious Diseases, Infection Prevention and Travel Medicine, Cantonal Hospital St Gallen, St Gallen, Switzerland; Division of Infectious Diseases & Hospital Epidemiology, University Hospital Basel, University of Basel, Basel, Switzerland; ALITHEA Genomics, Switzerland

## Abstract

The interaction between HIV-1 and the host immune system plays a crucial role in the natural control and progression of the infection. Previous studies have identified APOBEC3G, Tetherin, SAMHD1, and SERINC5 as HIV-1 host restriction factors, which are counteracted by the viral proteins Vif, Vpu, and Nef, respectively. The blood expression levels of some of these host proteins are correlated with HIV load, suggesting that interindividual differences in the spontaneous control of HIV infection might lead to the identification of novel HIV restriction factors.

Our study enrolled 150 participants from the Swiss HIV Cohort Study with human genome-wide genotyping data, pre-antiretroviral treatment peripheral blood mononuclear cells (PBMC) aliquots and HIV load measurements. Using BrB-seq, we quantified mRNA expression of all protein-coding genes and found significant associations between 792 genes and HIV load. Pathway analysis revealed that higher viral load associated with the upregulation of innate immune response, proteasome complex, mitochondrial and cell-cycle related activity, and with the downregulation of ribosomal transcripts and genes involved in cytokine-cytokine receptor interaction, including *IL4R*, *IL7R*, and *TCF7*. Mendelian Randomization confirmed the viral restriction activity of *TRABD2A* and identified new candidates as potential restriction factors. These findings provide novel insights into the host–virus interplay and suggest additional genes that may contribute to the natural control of HIV-1 infection.

## Introduction

Pathogen sensing is an essential first step in the host response to HIV-1 infection. This process provokes a cascade of immune reactions leading to the production of interferon and the consequent activation of interferon-responsive genes including restriction factors such as *APOBEC3G*, *SAMHD1* and Tetherin, which function to limit HIV replication and dissemination.

Different molecular mechanisms are involved in viral restriction that target various steps of the viral replication cycle(1, 2). Among the well known restriction factors, APOBEC3G is a single-strand cytidine deaminase that interferes with reverse transcription by inducing numerous deoxycytidine to deoxyuridine mutations in the negative strand of the HIV DNA primarily expressed as cDNA(3). SAMHD1(4), a cellular enzyme, hydrolyses nucleotide triphosphates (NTPs) to triphosphate and nucleosides, thereby reducing the amount of NTPs necessary for viral cDNA synthesis. Tetherin (encoded by the *BST2* gene) is an integral membrane protein that inhibits retrovirus infection by preventing the release of virus particles upon budding from infected cells(5). These restriction factors are counteracted by HIV accessory proteins: Vifmaps target APOBEC3G for proteasomal degradation by recruiting a Cul1-E3 ligase. Vpu promotes Tetherin degradation by recruiting SCFs ligase complexes(6).

More recently, SERINC5 and SERINC3 have emerged as constitutively expressed restriction factors that do not require activation by interferon(7, 8). Predominantly expressed in lymphocytes and monocytes, these proteins localize to the plasma membrane and act as a primary barrier against retroviral infection. We and others demonstrated that SERINC3 and SERINC5 are antagonized by the HIV-1 protein Nef. When expressed, Nef promotes the relocalization of SERINC3 and SERINC5 from the membrane to endosomal compartments with the involvement of clathrin-mediated trafficking. In absence of Nef, SERINC3 and SERINC5 are incorporated into virions and substantially inhibit viral infectivity(9, 10). While the exact restriction mechanism is not fully understood, the two proteins seem to act as lipid-flipping membrane transporters that disrupt viral membrane asymmetry, a function abrogated by Nef through interaction with intracellular loop 4(10). Importantly, despite the presence of Nef, it was shown that overexpressing *SERINC5* consistently impairs wild-type HIV-1(8) and the infectivity of other viruses(9), suggesting that inter-individual variations of expression of restriction factors (e.g., due to eQTLs) may confer a protective effect proportional to the respective expression levels.

Genome-wide association studies (GWAS)(11) have shown that host genetic variations have significant statistical associations with HIV-1 viral load (VL). Variants in the HLA region, especially HLA-B27 and HLA-B57, as well as CCR5Δ32, have been associated with better spontaneous control of HIV infection. A portion of the remaining variance may be a result of genetic variation in regulatory regions affecting immune gene expression, including that of restriction factors. Integration of transcriptomic data and GWAS findings provides a mean to uncover novel genes with antiviral capabilities and understand a link between variations in regulatory regions and HIV VL.

Building on these findings, we investigate here the transcriptomic landscape of individuals from the Swiss HIV Cohort Study (SHCS), stratified by set point VL in the absence of antiretroviral therapy. Our study is guided by the following hypotheses: i) viral control, as estimated by set point VL, correlates with the expression of known restriction factors such as *BST2*, *APOBEC* genes, *SERINC3/5* and others; ii) additional, previously uncharacterized restriction factors may exhibit similar correlations with viral load; iii) genetic variants (eQTLs) may upregulate restriction factors expression, thereby enhancing host defence; and iv) eQTLs may similarly modulate the expression of other unknown restriction factors with potential impact on HIV-1 control.

## Results

### 1. Transcriptomic profiling

From the SHCS, we enrolled 150 untreated people of European ancestry with HIV (PWH) based on set point VL, specifically including 75 individuals among the top quartile and 75 individuals among the bottom quartile of VL distribution (35 females, 115 males, mean age 39yo ranging from 24 to 68 – Supplementary Figure 1). Bulk RNA sequencing of peripheral blood mononuclear cells (PBMCs) was performed using the BrB-seq protocol(12) in two separate batches. Following initial quality control—retaining only samples with more than 6,000 expressed genes—a total of 144 samples were included in the mRNA expression analysis displaying between 1 and 15 million reads/sample and allowing the detection of 10’000 to 18’000 total genes/sample. (Supplementary Figure 2). Genes with no detectable expression across samples were excluded, resulting in 19,231 retained genes. Of these, 11,753 genes expressed in fewer than 10% of the samples were further filtered out, leaving 7,478 genes for downstream analyses. Raw gene counts were normalized to counts per million (CPM), log-transformed, and scaled.

[Data availability – Gene count matrices to be available in GEO NCBI XXX]

### 2. Associations between gene expression and HIV-1 viral load

For each gene, we explored the univariate interaction effect between normalized expression level and HIV-1 VL using a Log-Normal Generalized Linear Model. We also added the following covariates – sex, presence/absence of the HLA-B*27:05 and HLA-B*57:01 alleles, presence/absence of the CCR5 delta 32 deletion (rs333) and the 10 first principal components of the genotype matrix according to the formula:

**Equation 1.**

We calculated the effect sizes and their related p-values corrected with the Benjamini-Hochberg procedure for false discovery rate. The only covariate that was significantly associated with VL was sex (adjusted p-value = 0.02), as already observed(13), whereas a marginal contribution was provided by HLA-B*57:01 (0.058).

This analysis resulted in a list of 792 genes with significant associations between mRNA expression and HIV-1 VL. The top 20 positive and negative correlations are shown in Table 1 (full list in Supplementary Table 1). As an additional non-parametric measure of association, we calculated the Spearman correlation between gene expression and VL, identifying 586 of 792 genes with statistically significant correlations (FDR < 0.05).

**Table 1.**
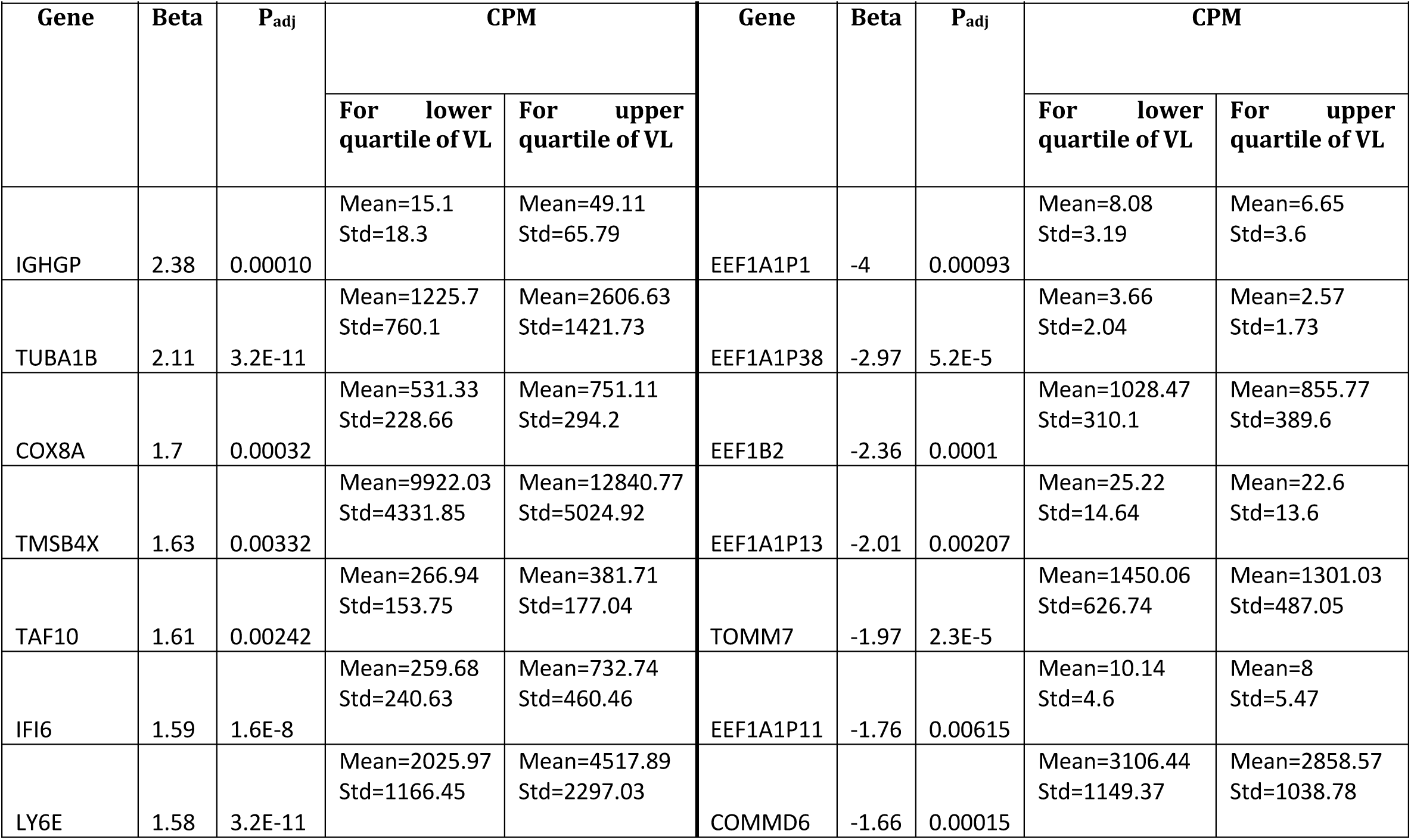
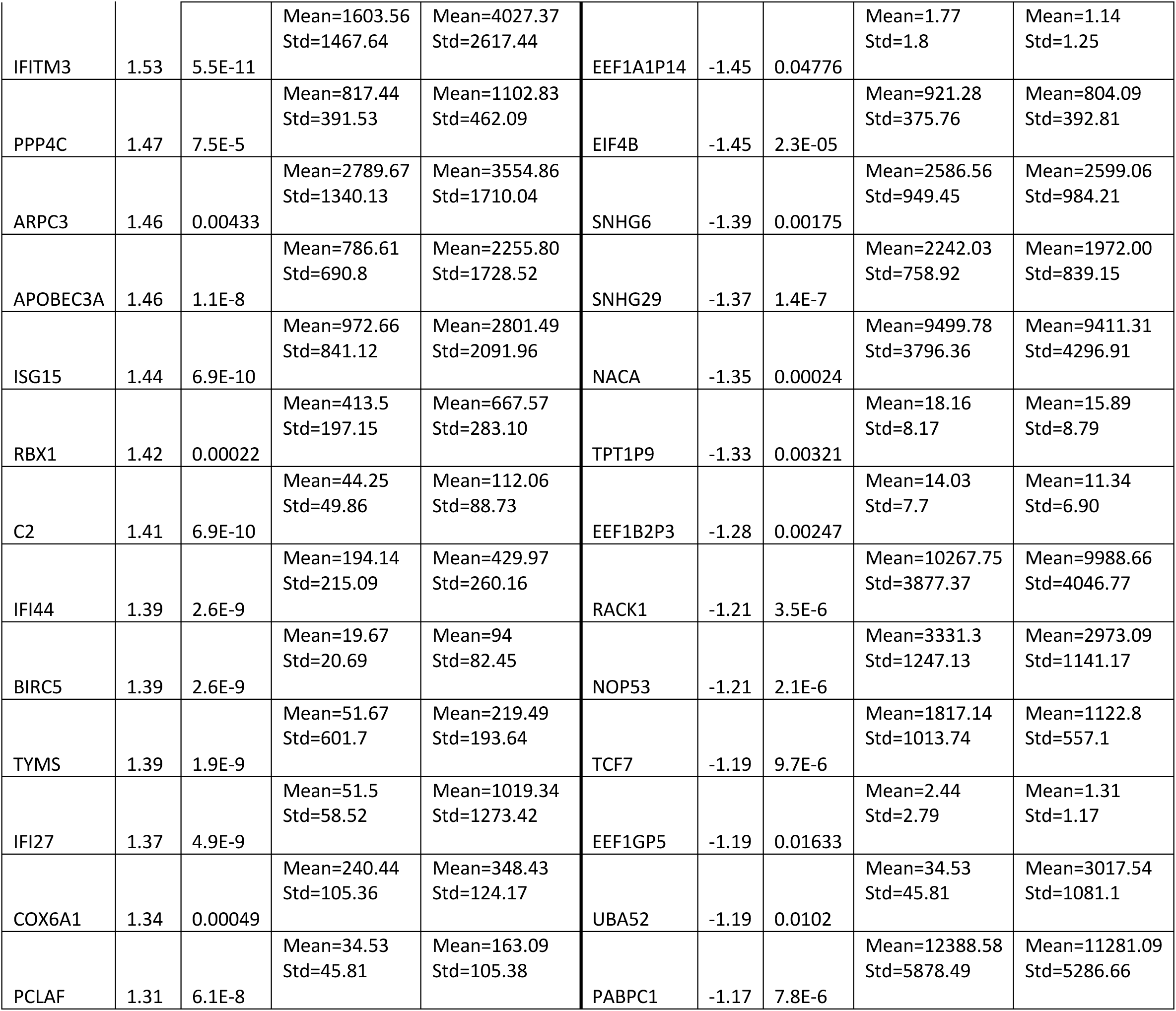
Top 20 genes with the strongest association between mRNA expression and HIV-1 viral load. We excluded ribosomal proteins from the list of negatively interacting genes.

Interestingly, the expression profiles of well-characterized HIV-1 restriction factors showed, overall, a positive correlation with viral load (VL) across individuals (Figure 1) such as *APOBEC3A* and *APOBEC3B*, members of the APOBEC3 family of cytidine deaminases(14). Another key factor positively correlated with VL is *BST2* (Tetherin), whose expression is induced by interferon. In general, out of the 792 genes, we detected VL association with 32 genes previously reported as restriction factors. Important interferon-induced genes include *IFITM2* and *IFITM3*, which restrict entry of various enveloped viruses(15) by interfering with membrane fusion via binding to the HIV-1 Env glycoprotein(16). Another antiviral factor targeting Env is *LGALS3BP*, which is active both intra-and extracellularly(17). Similarly, GBP5 restricts HIV-1 by blocking furin-mediated cleavage of Env, leading to reduced incorporation of functional glycoproteins(18). *OAS1*, *OAS2* and *OAS3* restrict HIV-1 by activating the RNase L pathway, leading to degradation of intracellular RNA(19). *RSAD2* (Viperin) has been reported to act against numerous viruses by inducing premature termination of RNA synthesis(20), but also to support sustained HIV-1 infection in monocyte-derived macrophages(21). Other interferon-induced factors include *ISG20*(22), *PLSCR1*(*23*), *SERPING1*(24) and others. Statistically significant associations of these genes appear to reflect an interferon-driven host immune response to elevated viral burden rather than intrinsic antiviral control. In contrast, *SERINC3* and *SERINC5*, constitutively expressed restriction factors not strongly induced by interferon, did not show significant correlation with VL. This lack of association in PBMCs may be due to their specific expression in macrophages, a cell type poorly represented in peripheral blood(7, 8, 25) or Nef-mediated viral counteraction mechanisms(26). Many genes previously shown to promote infection and participate in HIV-1 progression were positively correlated with VL. These included *CCR5*(27), *CFL1*(28), *LGALS1*(29), *LGALS3BP* (Galectin-3)(30), and *PPIB*(31). Heat shock proteins *HSP90B1*(32) and *HSPA4*(33) were also positively associated. Additional genes included members of the kinesin family (*KIF15*, *KIF18B*, *KIF2C*, *KIF5C*)(34), *ELOB*(35), and *STAT1*(36). Genes promoting viral transcription and reactivation were also enriched. These included *LGALS9*(37), *EZH2*(38), *BIRC5*(39) and *TYMS*(40).). Conversely, several host factors showed a negative association with HIV VL, including *RPLP1*(41), *GAS5*(42), *PRKRA*/*PACT*(43), *COMMD6*(44), *RCAN3*(45), *NCR3*(46), and *TCF7*(47). Pathway analysis indicated enrichment in high VL patients for antiviral defense (p = 3.4E-18) and response to interferon-beta (p=1.6E-4), cell division ( p= 1.1E-7), proteasome (p=2.4E-8), mitochondrion (p=1.6E-4) (Figure 2) Negatively associated pathways included cytoplasmic translation (p = 1E-127) and related ribosomal transcript production (4.4E-69) and cytokine–cytokine receptor interactions (3E-3) (Figure 3). The top 20 associated pathways are reported in Figure 4.

**Figure 1.**
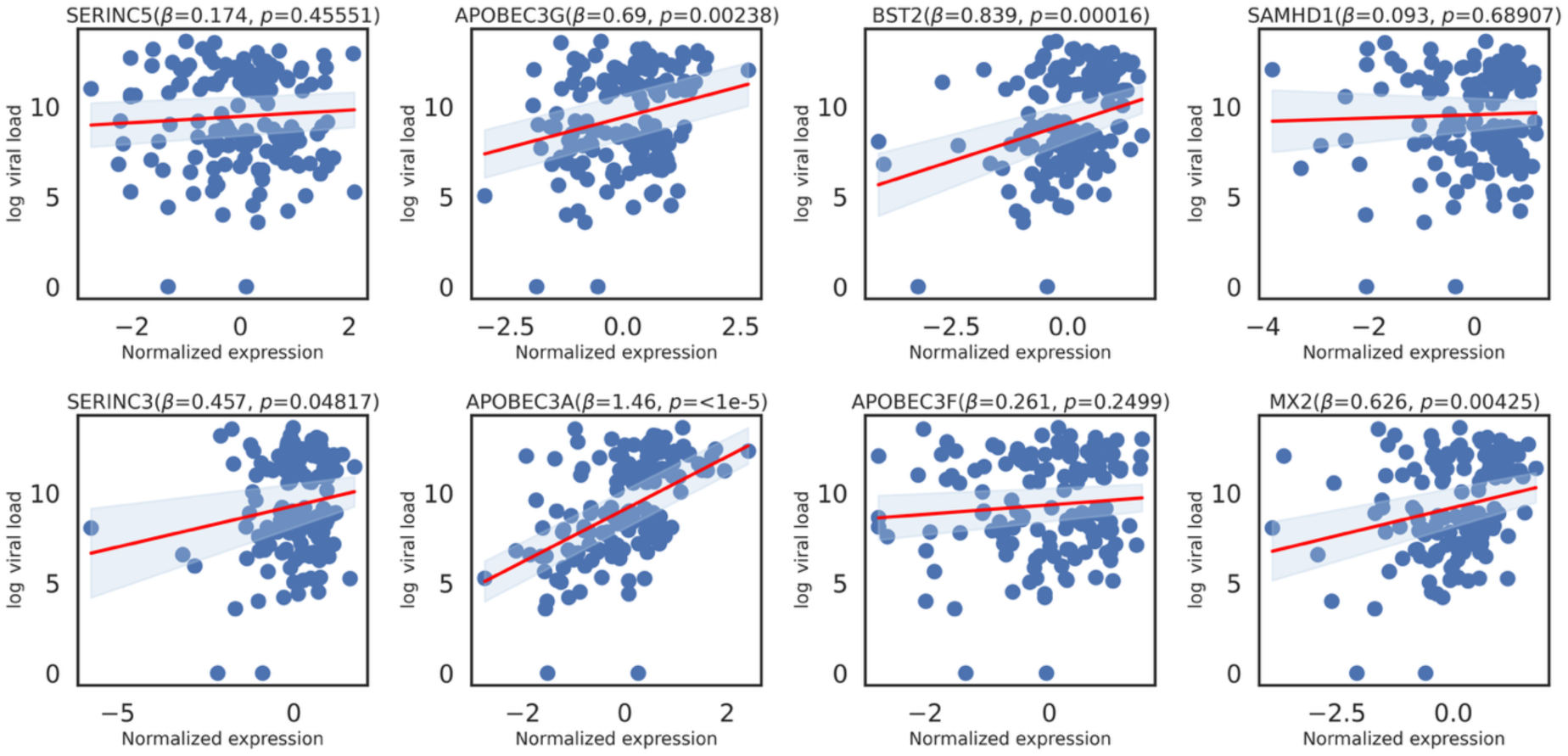
Correlations between mRNA expression levels of known HIV-1 restriction factors and HIV-1 viral load. Scatter plots show the relationship between normalized expression values (x-axis) of selected host restriction factors (SERINC5, APOBEC3G, BST2, SAMHD1, SERINC3, APOBEC3A, APOBEC3F, and MX2) and log-transformed plasma HIV-1 viral load (y-axis) in patient samples. Each blue dot represents an individual subject. Linear regression fits are shown in red, with 95% confidence intervals indicated by shaded areas. The slope (β) and significance level (p-value) for each correlation are reported in the panel titles. Significant positive correlations (e.g., APOBEC3G, BST2, APOBEC3A, and MX2) indicate that higher expression of these restriction factors is associated with higher viral load.

**Figure 2.**
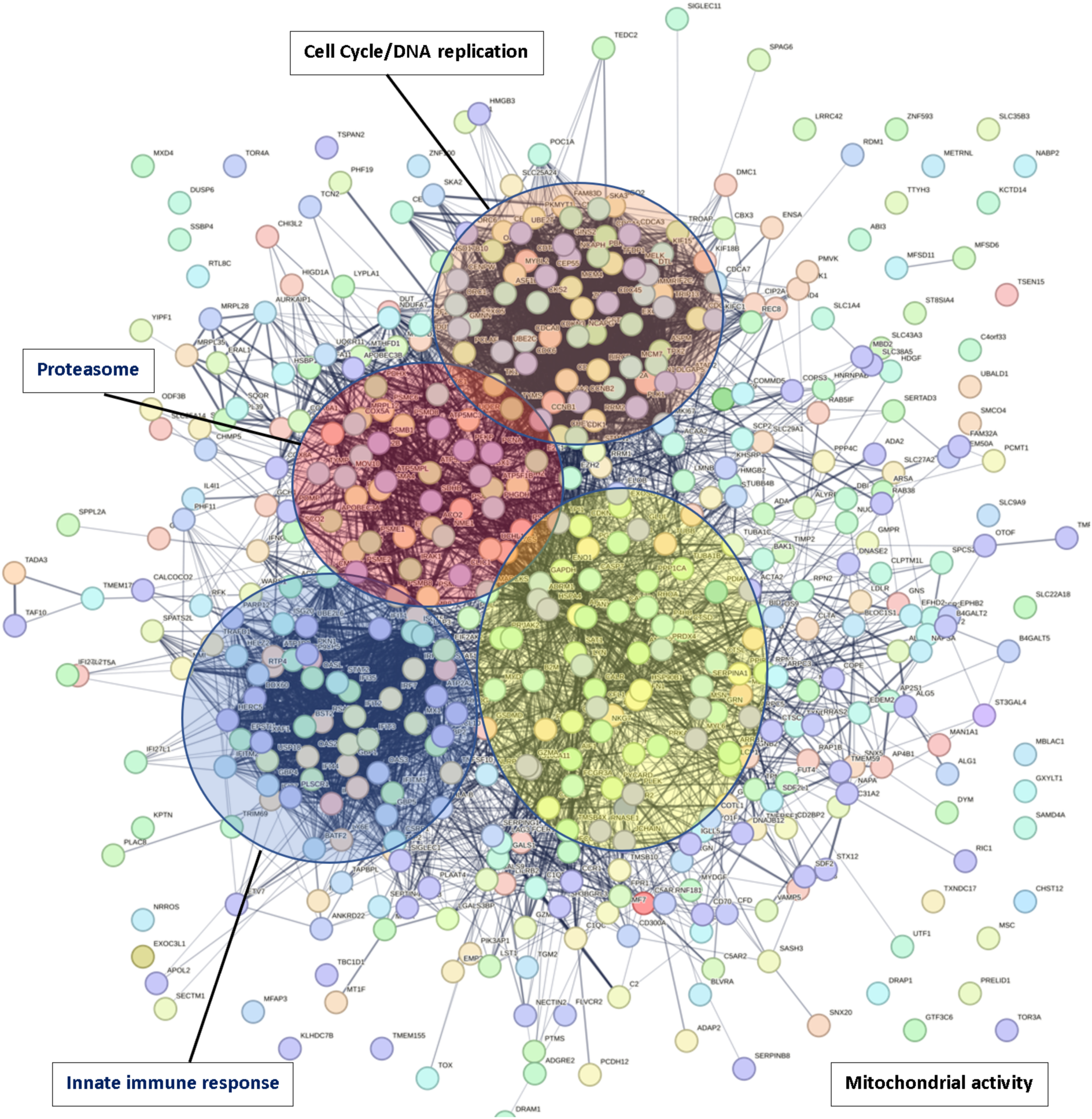
STRING database visualization of genes positively correlated with HIV viral load, and main related pathways. Protein–protein interaction (PPI) network generated using the STRING database of genes whose mRNA expression levels are positively correlated with HIV-1 viral load. Nodes represent individual proteins, and edges denote known or predicted functional associations. Colored circles highlight major functional clusters enriched among these genes, including pathways related to **cell cycle and DNA replication**, the **proteasome**, the **innate immune response**, and **mitochondrial activity**. The high connectivity within and between clusters suggests that HIV replication is associated with coordinated regulation of multiple host cellular processes.

**Figure 3.**
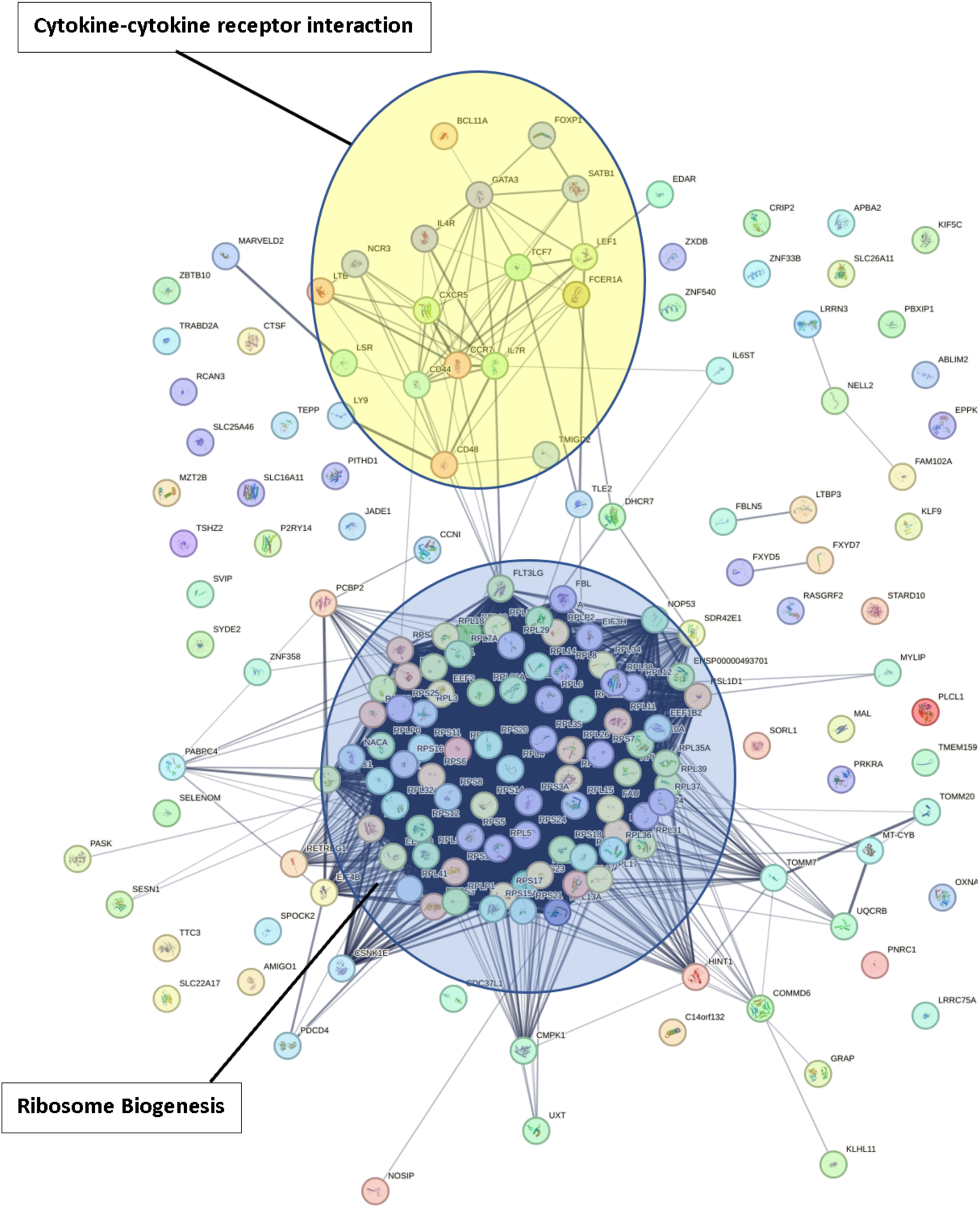
STRING database visualization of genes negatively correlated with HIV viral load and main related pathways. Protein–protein interaction (PPI) network of host genes whose mRNA expression levels are negatively correlated with HIV-1 viral load, constructed using the STRING database. Functional clustering highlights two major enriched pathways: **ribosome biogenesis** (dark blue cluster), representing a dense network of ribosomal proteins and assembly factors, and **cytokine–cytokine receptor interaction** (yellow cluster), consisting of immune signaling molecules and receptors. The observed negative correlations suggest that higher expression of ribosomal biogenesis and cytokine signaling genes is associated with lower HIV-1 viral load, implicating these pathways as potential host protective mechanisms against viral replication.

**Figure 4.**
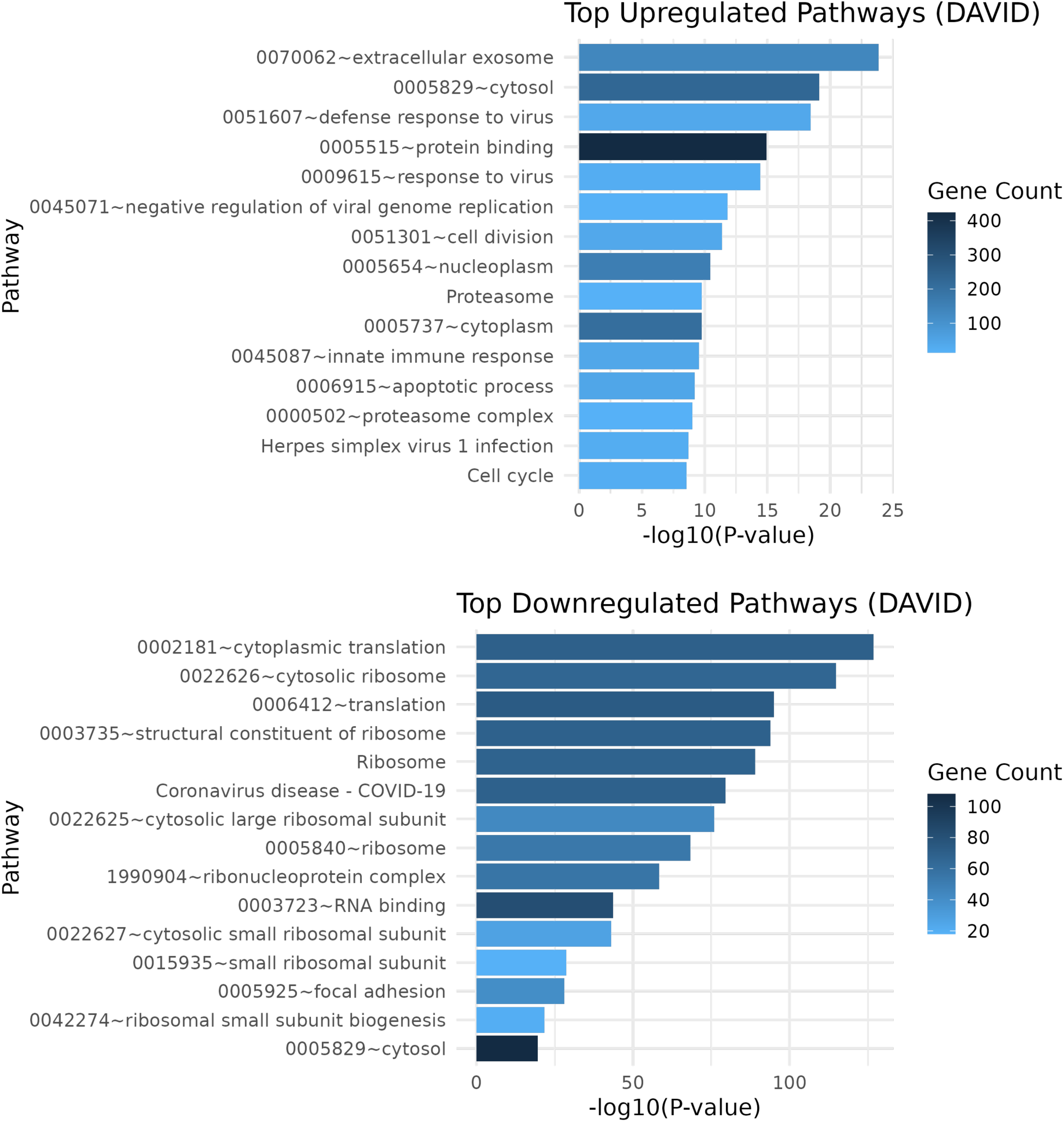
Differential pathway enrichment analysis of gene expression data using DAVID. Top: Significantly upregulated pathways, including extracellular exosome, cytosol, defence response to virus, protein binding, and negative regulation of viral genome replication, are shown with corresponding-log10(p-value) and gene counts. Bottom: Significantly downregulated pathways, predominantly related to cytoplasmic translation, ribosomal subunits, and RNA binding, are displayed with their-log10(p-value) and gene counts. Color intensity represents the number of genes contributing to each pathway.

### 4. Deconvolution of cellular composition

To interpret the results from the association analysis, we performed cell type deconvolution analysis of the gene expression of the PBMCs using EPIC(48). From the analysis of the whole transcriptome dataset, we observe that the majority of cells were CD8 T cells (up to 42%), B cells and NK cells whereas CD4+ T cells represented 10% of the PBMC population (Figure 5A). In addition, we compared the deconvolution of 25 samples with the lowest VL versus 25 samples with the highest VL (Figure 5B). As expected, low VL samples showed a significantly higher abundance of B cells and CD4 T cells compared to high VL samples, in which these populations were 70% depleted (log_2_ fold change=-1.76, p=0.029). Interestingly, the deconvolution of high VL samples showed a significant increase of neutrophils, supported by the positive association with VL of granulocyte-specific genes such as *CTSG*, *FPR1/2*, *SRGN*, and *GSDMD*. Indeed, even though granulocytes tend to be excluded from PBMC filtration, a subset of low-density neutrophils (LDN) and platelets might be retrieved in PBMC(49).

**Figure 5.**
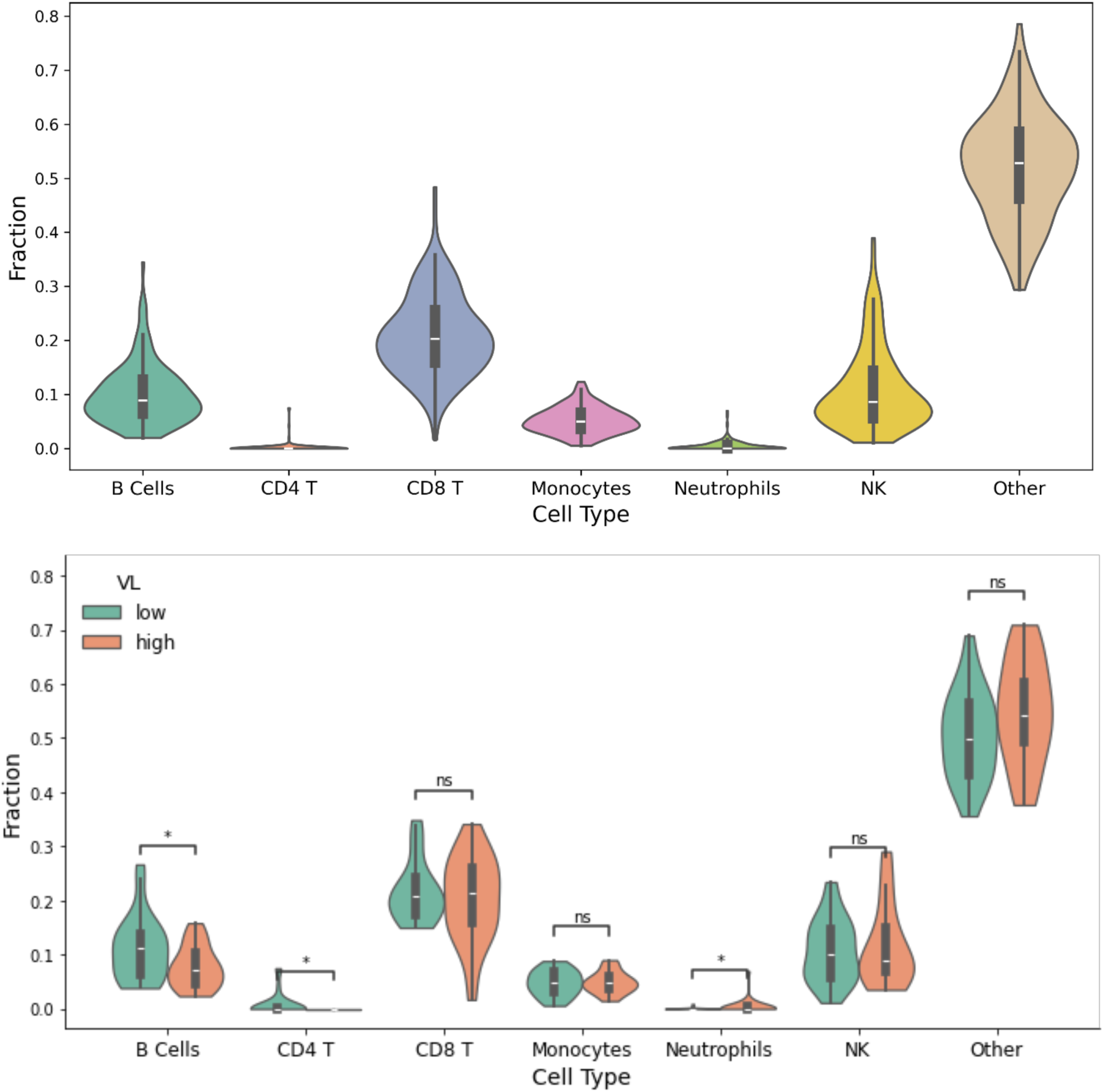
**Deconvolution of white blood cell populations**. A) Analysis of the full dataset reveals a depletion of CD4⁺ T cells and neutrophils. B) When comparing blood cell populations between PWH with low and high VL, B cells and CD4⁺ T cells are more abundant in the low VL group, whereas neutrophils are significantly more elevated in the high VL group (* P value < 0.05).

### 5. Mendelian randomization

We aimed to estimate the causal effect of the correlated genes on HIV-1 VL. A standard approach for assessing causality is Mendelian Randomization (MR), which leverages naturally occurring genetic variation as instrumental variables to infer the direction and magnitude of causal relationships between gene expression and disease phenotypes. To enable MR, we integrated expression quantitative trait loci (eQTL) data—linking genetic variants to gene expression levels—with GWAS summary statistics for HIV-1 VL. This integrative approach allowed us to investigate whether the observed correlations reflect true causal effects and to identify potential HIV restriction factor genes.

Using the Transcriptome Wide Mendelian Randomization (TWMR)(50, 51) approach applied to GWAS VL summary statistics and eQTLGen phase I data(52) for genes with observed associations between gene expression and VL, 146 genes were identified as causal, each supported by at least one eQTL. Given that for some of these genes there might be unobserved confounding, which violates key assumptions of MR, we additionally show the significance obtained with MRmix(53), MR-Egger(54), MR-presso(55), Mr-RAPS(56) and IVW(57) along with their robust(58)and penalized variations(59)(Figure 6 and Supplementary Table 2).

**Figure 6.**
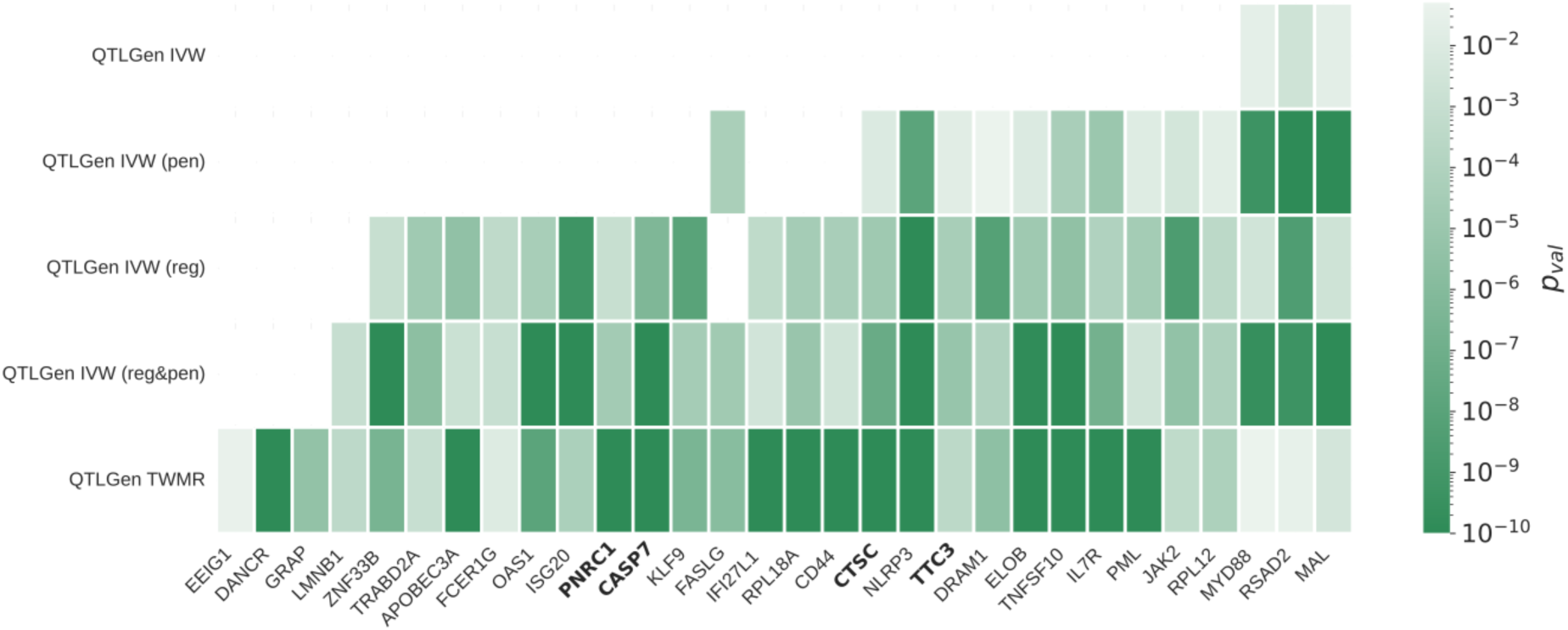
Identification of genes strongly correlated with VL. “Arkanoid”-style chart which represents genes with the strongest evidence of concordance across different types of IVW Mendelian Randomization methods, common GWAS summary on outcome (VL) and exposure (gene expression) dataset. Genes with no previous causal links to HIV-1 and its viral load are marked in bold. Visual representation of all investigated genes in Supplementary Figure 3.

Our analysis revealed that part of the genetic regulation of ribosomal protein genes (e.g., *RPL12* (p=7.1E-05, supporting MR methods: 9/23), *RPL17* (p<1×10⁻²², 4/13), *RPL18A* (p<1×10⁻²², 5/13), *RPL23A* (p<1×10⁻²², 3/11), and others; Supplementary Table 2) is causally and negatively associated with VL, supporting the hypothesis that HIV-infected cells downregulate ribosomal pathways to mitigate viral replication(60). In line with this, several genes involved in peptide chain elongation (e.g., *EEF1B2* (p<1×10⁻²², 1/12), *ELOB* (p<1×10⁻²², 6/12) also showed causal associations, suggesting that host genetic variation may influence interactions between HIV-1 and the reverse transcription complex(61).

Among the negatively correlated genes with a significant causal effect on VL, *TRABD2A* (p=0.003, 11/20) stands out as a recently identified HIV-1 restriction factor. Another strong signal was *PNRC1* (p< 1×10⁻²², 3/13), a nuclear receptor coactivator involved in mRNA decay and transcriptional regulation. We also identified *IL7R* (p<1×10⁻²², 6/12), encoding the interleukin-7 receptor. Other notable genes with previously observed interaction with HIV-1 are *EEIG1* (p=0.0375, 2/9), the early estrogen-induced gene 1; *MAL* (p=0.0037, 6/9), a myelin and lymphocyte protein associated with lipid raft signaling in immune cells; the long non-coding RNA DANCR (p<1×10⁻²², 4/12) expressed during the early stages of HIV-1 infection. Surprisingly, *CD44* (p<1×10⁻²², 4/13) is part of this group of genes despite being tagged as a host factor incorporated in virions to promote hyaluronan-mediated infection of Fibroblastic reticular cells(62).

Finally, we detected associations with genes not previously known as HIV-1 interactors, such as *TTC3* (p=0.0003, 8/20), *ZNF33B* (p=2.6×10⁻⁷, 4/13), *KLF9* (p=3e-7, 6/13), *GRAP* (p<4e-6, 3/13). These are intriguing candidates for further functional validation and may represent novel host factors influencing HIV-1 replication or latency.

Among the positively correlated genes, several apoptosis-related factors emerged, including *FASLG* (p=1.4×10⁻⁶, 3/12), *TNFSF10* (p<1×10⁻²², 6/12), *CASP7* (p<1×10⁻²², 6/23), *CTSC* (p<1×10⁻²², 7/19), *LMNB1* (p=3.6×10⁻⁴, 3/13), and *DRAM1* (p=2.6×10⁻⁶, 7/23).

We also identified a set of innate immunity–related genes—many of which are known interferon-stimulated genes (ISGs)—with strong causal links to HIV-1 VL (GO enrichment, adjusted p=2×10⁻³). These include well-established HIV restriction or response factors such as *APOBEC3A* (p=1.1×10⁻¹², 8/24), *ISG20* (p=5.7×10⁻⁵, 7/20), *RSAD2*/viperin (p=0.03, 8/12), *PML* (p<1×10⁻²², 7/13), *JAK2* (p=4.7×10⁻⁴, 7/13), *MYD88* (p=0.0498, 6/9), *OAS1* (p=1.2×10⁻⁸, 6/20), *FCER1G* (p=0.01, 5/23), *NLRP3* (p<1×10⁻²², 6/23), and *IFI27L1* (p<1×10⁻²², 6/23).

## Discussion

We explored the interaction between PBMC gene expression and HIV-1 set point VL as a proxy for spontaneous viral control in 144 PWH. Using a Log-Normal Generalized Linear Model, we identified 792 genes whose expression levels were significantly positively or negatively associated with VL. Together, these results highlight a complex interplay between host restriction factors, interferon-driven immune responses, and viral countermeasures in shaping HIV-1 dynamics. Interferon-induced antiviral genes (e.g., *APOBEC3* family, *BST2*, *IFITM*s, *OAS* family) correlated positively with VL, likely reflecting virally induced immune activation rather than direct control of replication. Additionally, our cell type deconvolution revealed a depletion of B and CD4⁺ T cells and an enrichment of neutrophils in high VL samples. We hypothesize that the loss of B and CD4⁺ T cells is partly compensated by a shift toward innate, but non-specific, neutrophil-driven responses, where low-density neutrophils suppress adaptive immunity against HIV-1 through PD-L1/PD-1 and expand under high viral burden(63).

The enrichment of host dependency factors (*CCR5*, *CFL1*, *LGALS* family, *PPIB*, kinesins) among positively correlated genes reinforces their central role in HIV-1 replication and progression. Associations with transcriptional regulators (*EZH2*, *STAT1*, *LGALS9*) and heat shock proteins suggest mechanisms that facilitate viral reactivation and latency maintenance. Pathway-level enrichments in translation, NF-κB/NFAT signaling, and proteostasis further point to regulatory nodes that could be targeted therapeutically.

While no significant association was observed for *SERINC3* and *SERINC5*, we found several negatively associated genes (*RPLP1*, *GAS5*, *PRKRA*, *COMMD6*, *RCAN3*, *TCF7*) recently characterized as important regulators of T-cell metabolism and viral control. For instance, TCF7 mRNA expression in PBMCs has been reported to be lower in rapid progressors compared to long-term non-progressors and healthy controls(64), and *TCF7* knockdown disrupts mitochondrial function and reduces CD8+ T-cell proliferative capacity(65), highlighting the central role of metabolic regulation in TCF7-mediated antiviral control.

We also observed a negative association between VL and the expression of eukaryotic translation initiation and elongation factors (*EEF1B2*, *EEF1D*, *EEF2*, *EIF3H*, *EIF4B* and pseudogenes *EEF1A1P1*, *EEF1A1P13*), of which *EIF3H*, *EEF1D* and *EEF2* have already been reported as exploited by HIV-1 for viral protein synthesis(66–68). Consistent with prior findings, several ribosomal transcripts were depleted at high VL, suggesting translational repression as a host restriction mechanism. Additional negatively associated causal genes included *TRABD2A*, which inhibits virion assembly by degrading the Gag polyprotein, *PNRC1*, a nuclear receptor coactivator involved in mRNA decay, *IL7R*, *MAL*, *DANCR*, and *GRAP*. Positive correlations were observed for genes involved in viral entry, replication, latency, and apoptosis, including *CCR5*, *CFL1*, *LGALS1*/*3BP*, *PPIB*, heat shock proteins (*HSP90B1*, *HSPA4*), *LGALS9*, *EZH2*, *BIRC5*, *TYMS*, and *DRAM1*. Notably, *DRAM1* is upregulated by HIV via p53 and may contribute to the elimination of infected cells(69), highlighting the interplay between host apoptotic pathways and viral replication.

Gene ontology and pathway analyses revealed a downregulation of ribosomal biogenesis and cytokine-cytokine receptor interactions at high VL, whereas interferon-stimulated genes, proteasome activity, cell-cycle/DNA replication, and mitochondrial pathways were upregulated. These results suggest that HIV-1 induces host transcriptional programs to enhance replication while simultaneously activating antiviral defense mechanisms. Mendelian Randomization further clarified causal relationships, identifying *IFI35* and components of the MHC Class I peptide-loading complex as modulators of VL, as well as members of the *OAS* family working with *EIF2AK2* to restrict infection.

Our study also identified the TNFSF10 (TRAIL) signaling pathway as a potential mediator of HIV-1–induced CD4⁺ T cell depletion. Previous work in humanized (NRG-hu HSC) mice demonstrated that blockade of TRAIL prevented HIV-1–induced CD4⁺ T cell loss in vivo(70). Here, we observed transcriptomic signatures consistent with TRAIL+ innate immune cells, supporting the role of TRAIL-mediated apoptosis in shaping CD4⁺ T cell dynamics during infection.

The identification of *TTC3*, *ZNF33B*, *KLF9*, and *GRAP* as genes associated with HIV-1 VL is particularly intriguing, as none of these factors have previously been characterized as HIV-1 interactors. *TTC3* encodes a tetratricopeptide repeat protein with E3 ubiquitin ligase activity, involved in protein ubiquitination and degradation of phosphorylated Akt(71). Since it has already been shown that PI3K/Akt survival pathway is triggered by HIV to prevent apoptosis of HIV-infected cells(72), *TTC3* may counteract this effect by promoting Akt degradation, thereby influencing the balance between survival and death of infected cells and potentially impacting reservoir persistence. *ZNF33B*, an uncharacterized zinc finger transcription factor, could influence HIV-1 by modulating host chromatin or viral promoter activity, as other zinc finger proteins (e.g., ZNF restriction factors) have been implicated in HIV-1 viral replication(73).

*KLF9*, a ubiquitously expressed member of the Sp1 C2H2-type zinc finger family of transcription factors, has roles in T-cell differentiation(74) and stress responses(75). It was previously observed as a possible activator of HIV-1 LTR promoter via GC-box binding(76) therefore its negative association with VL is unexpected. However, as other member of the *KLF* family, *KLF9* might also act a transcriptional repressor by competing with SP1(77, 78). Finally, *GRAP* encodes a lymphoid-specific adaptor protein in the Ras/MAPK pathway, a signaling cascade activated by T-cell receptor stimulation(79) and known to facilitate HIV-1 replication and reactivation from latency(80).

In summary, our integrative analysis combining PBMC transcriptomics and host genotypes provides an improved overview of the host factors associated with HIV-1 VL control. We confirm the importance of interferon-dependent restriction factors, translational repression, apoptotic pathways, and T-cell metabolic regulators, while also highlighting novel candidate genes and pathways, including TRAIL, that may influence viral control. These findings provide new insights to further understand virus-host interactions and identify novel putative targets for validation in future functional studies and for potential therapeutic interventions.

## Methods

### Study participants, samples and data

The SHCS (www.shcs.ch) is an ongoing, nationally representative cohort study of PWH. Clinical, behavioral, and laboratory data are collected at study registration and biannually thereafter. All centers’ local ethical committees approved the cohort study, and all participants provided written informed consent, including for human genetic testing.

For this study, we selected 150 SHCS participants of European ancestry, who had genome-wide genotyping data from previous studies and a PBMC aliquot available before antiretroviral treatment. PWH were selected based on their VL at set point (75 individuals among the top quartile; 75 individuals among the bottom quartile of the VL distribution). The following clinical and laboratory parameters were extracted from the SHCS database and included in the analysis as stratifying factors and/or covariates: sex at birth, age at time of PBMC sampling, presence/absence of HLA-B*27:05, HLA-B*57:01 and the CCR5 delta 32 deletion (rs333), setpoint VL, number of CD4+ T cells and VL at time of PBMC sampling.

### DNA genotyping

Genotyping was performed in the context of previous SHCS projects using genotyping arrays from Illumina and DNA extracted from PBMCs. For each genotyping batch, samples and single nucleotide polymorphisms (SNPs) were removed if the missingness exceeded 10% or the minor allele frequency deviated more than 20% from the 1000 Genomes Project reference. Missing genotypes were phased with EAGLE2(81) and imputed using positional Burrows-Wheeler transformation (PBWT)(81), at the Sanger Imputation Service, using the 1000 Genomes Project Phase 3 panel as reference. Only high-quality SNPs with an imputation information score (INFO > 0.8) were retained following the imputation, after which the genotyping batches were combined. Principal components and, separately, population structure together with the 1000 Genomes reference panel was calculated using PLINK2 (v2.00a3LM). KING (v2.1.3) was used to ensure that no duplicate or cryptically related samples were included in the final dataset.

### RNA extraction and sequencing

PBMC samples (∼10^6^ cells) were stored at-80°C. They were thawed in a water bath at 37°C under manual agitation and rapidly resuspended and washed in 10ml warm PBS (37°C). Tubes were centrifuged 5min at 1100rpm, PBS removed and a second washing step with warm PBS was performed. Pellets were resuspended immediately in 350µl RLT plus with β-mercaptoethanol (1:100) for cells lysis and processed for RNA extraction according to manufacturer’s instructions (RNeasy Plus Mini Kit, Qiagen Cat.No./ID:74134). Elution volumes were 18µl and final concentrations were mostly between 15 and 50 ng/µl. From this step, all manipulations were done on ice. RNA quality (RIN: RNA Integrity Number) was analyzed using a Tapestation instrument and the RNA ScreenTape system kit (Agilent, ref. 5067-5576 and 5067-5577). RNA quantification was performed using the Qubit^TM^ system and the RNA Quantification high sensitivity kit (ref.: Q32852)

To perform RNA sequencing, we applied the BrB-seq protocol (Alithea Genomics), which enables the simultaneous generation of RNA-seq libraries through high sample multiplexing and quantification of gene expression from 3’cDNA libraries(12). The libraries were sequenced with an average coverage of 5Mio reads at the Gene Expression core facility of the EPFL on Illumina sequencers.

### Gene expression and eQTL analyses

The calculation of gene expression matrices was obtained through a proprietary pipeline provided by Alithea Genomics. Preprocessing of raw gene counts was done using Scanpy (82) toolkit and in-house developed python scripts, which perform normalization of counts, log-transform and batch correction. We used DAVID Functional Classification Tool(83) for gene ontology analysis, including clustering and annotation charts. We used STRING(84) with standard settings for gene interaction network analysis.

### Mendelian randomization

Mendelian Randomization is a convenient way to elucidate the causal relationships between an exposure and an outcome, provided that human genetic variants can be used as instrumental variables to “randomize” the exposure. We utilized different Mendelian Randomization approaches to elucidate the causal impact of genetic variation on gene expression and thereby assess interactions between HIV-1 viral load and gene expression attributed to genetic variation in the host rather than infection itself. We applied classic inverse-variance weighting IVW along with regularized and penalized variations, and TWMR with gene expression is set as exposure, HIV-1 VL as outcome and SNPs as instrumental variables. Additional methods were used to support TWMR primary estimation. Outcome effects of QTLs on VL come from a single GWAS summary dataset, for exposure QTL effect on gene expression we utilize QTLGen dataset as primary source of estimations for whole blood.

For each MR method except TWMR we employ a set of independent QTLs with LD R^2 < 0.1 with a nominally significant (p-value < 0.05) effect both on gene expression and viral load. As TWMR while computing gene causal effect considers also effects of genes, regulated by the same QTLs in order to account for possible pleiotropy, in context of this method for each focal gene we first assemble set of independent QTLs(LD R^2 < 0.1) for a focal gene, then add additional genes regulated by these QTLs disregarding of their possible effect on outcome, then extend initial QTL set with all independent QTLs which affect both focal and adjacent genes. In order to avoid as much pleiotropy as possible, we retain only independent QTLs by picking only QTLs with the lowest standard error from each LD block with R^2 > 0.1.

### Financial support

This study has been financed within the framework of the Swiss HIV Cohort Study, supported by the Swiss National Science Foundation (grant #33FI-0_229621), by SHCS project #812 and by the SHCS research foundation. The data are gathered by the five Swiss University Hospitals, two Cantonal Hospitals, affiliated hospitals and private physicians.

### Members of the Swiss HIV Cohort Study

Abela IA, Aebi-Popp K, Anagnostopoulos A, Bernasconi E, Braun DL, Bucher HC, Calmy A, Cavassini M (Chairman of the Clinical and Laboratory Committee), Ciuffi A, Dollenmaier G, Egger M, Elzi L, Fehr JS, Fellay J, Frigerio Malossa S, Fux CA, Günthard HF, Hachfeld A, Haerry DHU (deputy of “Positive Council”), Hasse B, Hirsch HH, Hoffmann M, Hösli I, Huber M, Jackson-Perry D (patient representatives), Kahlert CR, Kaufmann D, Keiser O, Klimkait T, Kouyos RD, Kovari H, Kusejko K (Head of Data Centre), Labhardt ND, Leuzinger K, Martinez de Tejada B, Marzolini C, Metzner KJ, Müller N, Nemeth J, Nicca D, Notter J, Paioni P (Chairman of the Mother & Child Substudy), Pantaleo G, Perreau M, Rauch A (President of the SHCS), Salazar-Vizcaya LP, Schmid P, Segeral O, Speck RF, Stöckle M, Surial B, Tarr PE, Trkola A, Wandeler G (Chairman of the Scientific Board), Weisser M, Yerly S.

## Supplementary Figures

**Supplementary Figure 1.**
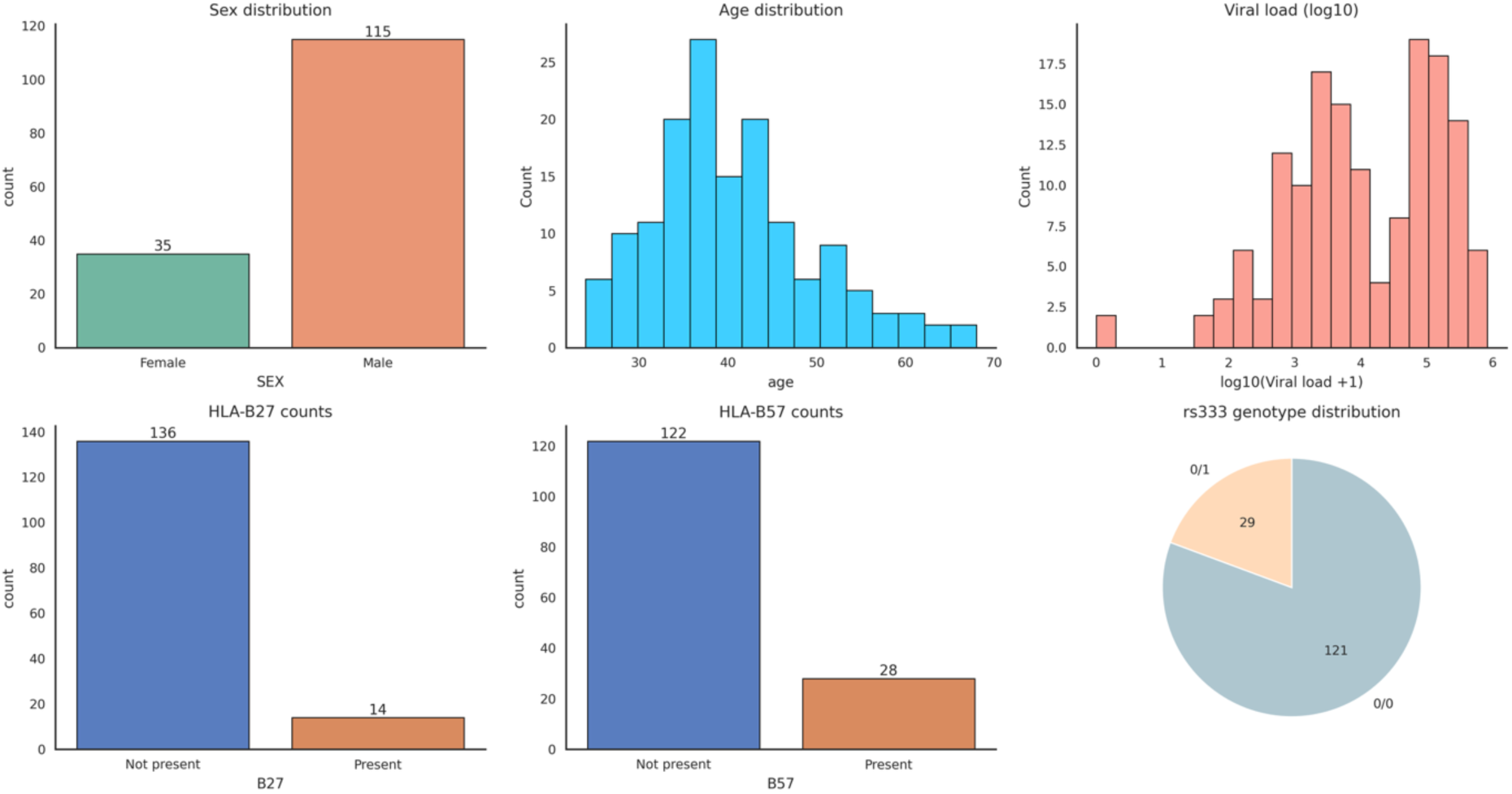
Distributions of sample features. A) Number of male and female samples, B) distribution of patients in samples, C) Distribution of log10 of viral load in samples D) Number of samples with and without HLA-B27 genotype E) Number of samples with and without HLA-B57 genotype F) Number of rs333 genetic variant carriers with wild type denoted as 0/0, heterozygous – 0/1

**Supplementary Figure 2.**
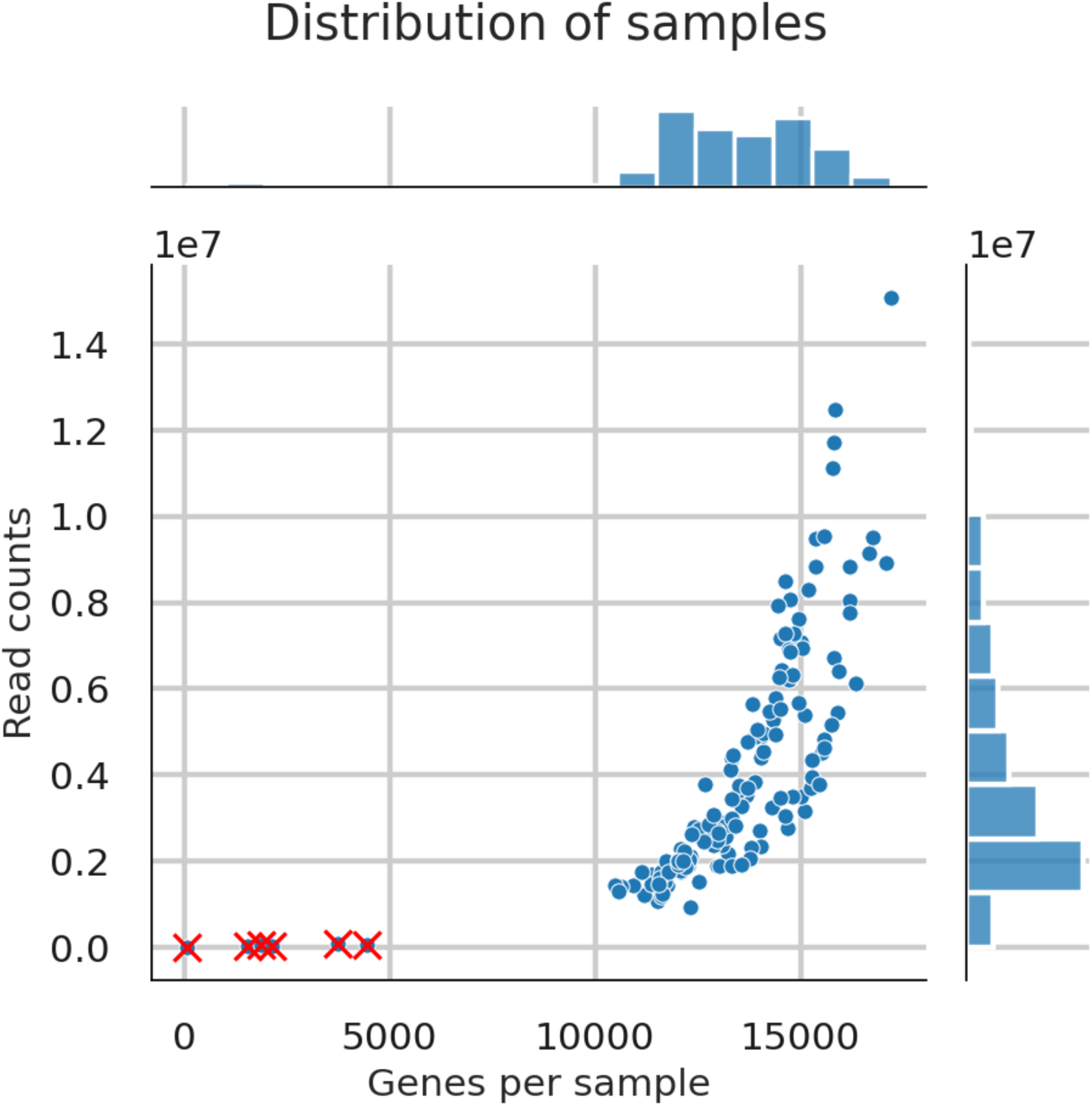
Distribution of read counts per number of expressed genes in 150 samples. Each dot represents an individual sample. Marginal histograms display the distribution of expressed genes (top panel) and total read counts (right panel), highlighting the overall variability across the dataset. Crossed out dots represent samples excluded from analysis due to low sequencing quality.

