## Supplementary figures and images for "Interdependent Dynamics of mRNA Expression and HIV-1 Viral Load: Insights from Transcriptomics and Mendelian Randomization"

### Supplementary Figure 3

#### MR p-values and gene expression results

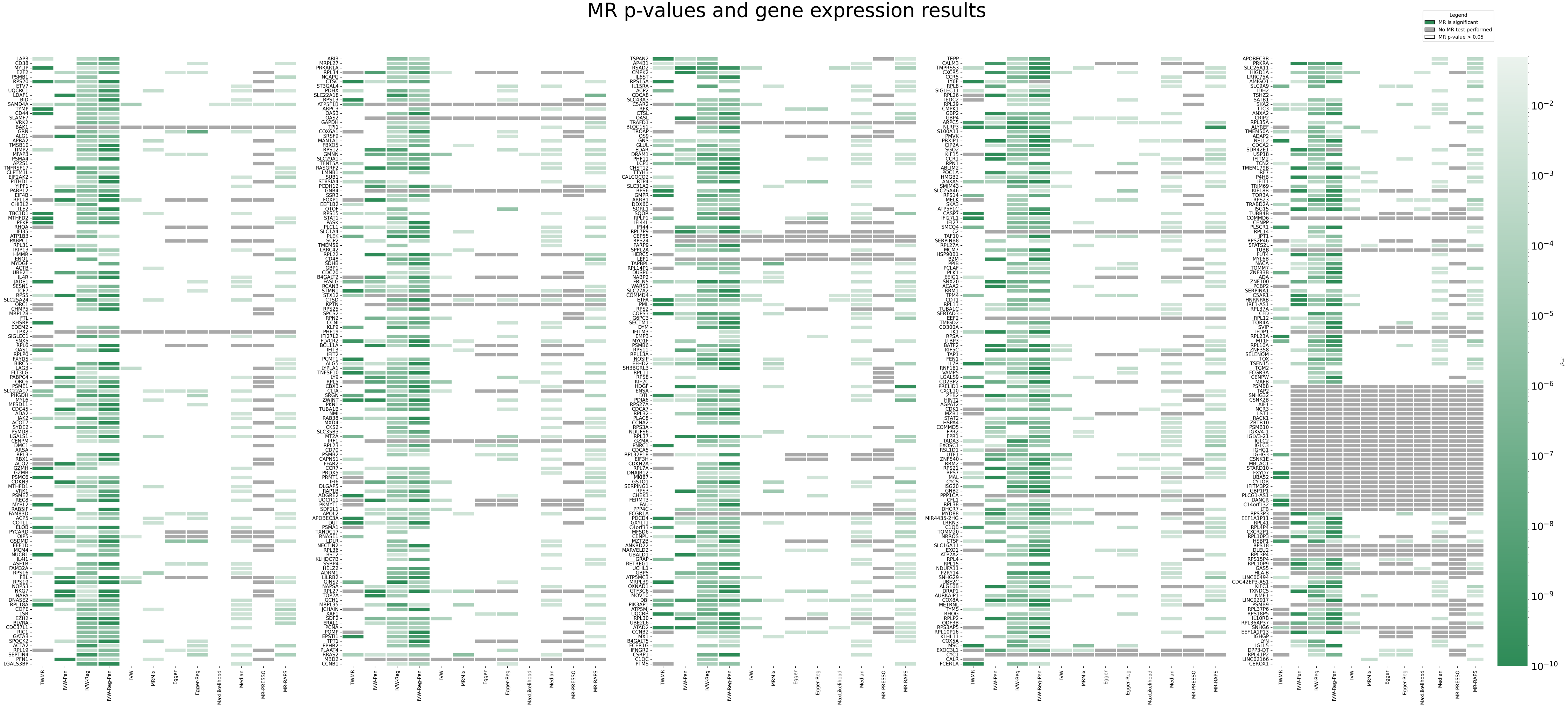
